# Pre-treatment naïve T cells are associated with severe irAE following PD-1/CTLA4 checkpoint blockade for melanoma

**DOI:** 10.1101/2025.10.30.685578

**Authors:** Kathryne E. Marks, Alice Horisberger, Mehreen Elahee, Ifeoluwakiisi Adejoorin, Nilasha Ghosh, Michael A. Postow, Laura Donlin, Anne R. Bass, Deepak A. Rao

## Abstract

Immune checkpoint inhibitors (ICIs) such as anti-PD-1 and anti-CTLA-4 antibodies are used to induce an immune response against many types of tumors. However, ICIs often also induce autoimmune responses, referred to as immune-related adverse events (irAEs), which occur unpredictably and at varying levels of severity in ICI-treated patients. The immunologic factors that predispose patients to the development of severe irAE are largely unclear. Here, we utilized high dimensional mass cytometry immunophenotyping of longitudinal blood samples from patients with metastatic melanoma treated with combination anti-PD-1/CTLA4 ICI therapy in the context of a clinical trial to characterize alterations in immune profiles induced by combination ICI therapy and to identify immune features associated with development of severe irAEs. Deep T cell profiling highlighted that ICI therapy induces prominent expansions of activated, CD38^hi^ CD4^+^ and CD8^+^ T cells, which are frequently bound by the therapeutic anti-PD-1 antibody, as well as substantial changes in regulatory T cell phenotypes. However, neither the baseline frequency nor the extent of expansion of these cell populations was associated with development of severe irAEs. Rather, single cell-association testing revealed naïve CD4^+^ T cell abundance pre-treatment as significantly associated with the development of severe irAEs. Biaxial gating of naïve CD4^+^ T cells confirmed a significant positive association of naïve CD4^+^ T cell proportion and development of a severe irAE and with the number of irAEs developed in this cohort. Results from this broad profiling study indicate the abundance of naïve CD4^+^ T cells as a predictive feature for the development of severe irAEs following combination anti-PD-1/CTLA4 ICI therapy.

## Introduction

Immune checkpoint inhibitors (ICIs) have significantly advanced cancer treatment since their initial approval over a decade ago and contribute to lasting survival (1-3). ICI therapy with antibodies that block PD-1, PD-L1, CTLA-4, or LAG-3 have led to improved responses against cancers of skin, lung, kidney and other tissues (4), and treatment with a combination of checkpoint blockade treatments can increase their efficacy (5). ICI therapies release T cells from inhibitory signals and activate them to recognize and kill tumor cells. However, ICI therapies can also cause systemic changes in T cell function, in addition to their intended effects on tumor-infiltrating T cells. Changes to circulating T cell populations following combination of ICI therapies, such as anti-PD-1 together with anti-CTLA-4, can be distinct from treatment with either monotherapy (6-8). The extent of changes to well-defined subsets of circulating immune cell populations following combination PD-1/CTLA4 blockade remains incompletely defined.

Systemic or extra-tumor T cell activation by ICI therapies can induce immune-related adverse events (irAEs) (9, 10). irAEs range in severity, timing, and organ system affected, but are a frequent result of ICI therapies, occurring in up to 90% of people receiving combination anti-PD-1/CTLA-4 blockade therapy (11, 12). More than half of irAEs resulting from this combination are grade 3 or 4 and can necessitate hospitalization and cessation of therapy (11). Common irAEs include colitis, rashes, hepatitis, and arthritis among others (10, 13). While myositis, and myocarditis are rarer irAEs, they are often severe and can result in mortality (10).

Specific clinical factors have been associated with increased risk of irAE or more severe irAEs in some studies, including high BMI, smoking, and male sex; however, these associations have not been consistently observed (14-17). Younger age has also been associated with more severe irAEs in some but not all studies (18, 19). Identification of cells in circulation that are associated with type or severity of irAE before or early in treatment would enable better prediction of irAE development. Studies of cellular profiles have yielded varied results regarding associations of T cell phenotypes associated with irAEs (20-22). CD4^+^ T cells, particularly memory CD4^+^ T cells and more TCR diversity were reported in an observational study as associated with future irAE severity in a cohort of both anti-PD-1 alone and combination therapy-treated patients (22). However, a separate observational study found that naïve CD4^+^ T cells were present at higher levels in patients with higher severity irAEs following either anti-PD-1, anti-CTLA-4, or combination anti-CTLA4/anti-PD-1 therapy (21). Another observational study utilizing flow cytometry found no differences in naïve CD4^+^ or CD8^+^ T cells in patients treated with ICI therapy with an irAE vs without any irAEs (20), thus, substantial uncertainty remains about T cell phenotypes predictive of irAE development.

Utilizing PBMCs collected from well characterized patients through the Adaptively Dosed ImmunoTherapy Trial (ADAPT-IT) (5), we aimed to determine the specific effects of anti-PD-1/CTLA-4 combination therapy on immune cell populations in blood and to determine associations of blood immune cell features with irAE severity in the setting of this clinical trial. Previous analysis of patients in the ADAPT-IT trial revealed an increase in proliferating T cells after 1 dose of combination therapy. Here, we extend the analysis of patients in this trial by using high-dimensional mass cytometry profiling of circulating T cells to identify cellular features in the baseline, pre-treatment samples associated with irAE development (23). We show dramatic expansion of activated and proliferating T cells that are directly acted upon by anti-PD-1 therapy. Notably, this dramatic cellular activation is not associated with development of irAEs. Rather, our broad immune profiling approach demonstrated that the level of naïve T cells, especially naïve CD4^+^ T cells, before ICI therapy is significantly associated with development of severe irAEs following combination anti-PD-1/CTLA-4 therapy in this clinical trial setting.

## Results

### Patient characteristics

We obtained PBMCs from patients with advanced melanoma enrolled in the ADAPT-IT trial (5). Patients in this multicenter, single-arm, phase II clinical trial were treated with 2 doses of both nivolumab (1 mg/kg), an anti-PD-1 therapy, and ipilimumab (3 mg/kg), an anti-CTLA-4 therapy, followed either by nivolumab monotherapy every 2 or 4 weeks or 2 more doses of combination therapy followed by nivolumab monotherapy every 2 or 4 weeks. Patients in this trial had no prior exposure to ICI-therapy and no active autoimmune disease at the time of participation. The demographics of analyzed patients are summarized in Table 1 and Supplementary Table 1 and detailed in Supplementary Table 2. Among the overall cohort, the mean age was 60 years and the cohort was 59% male.

**Table 1.**
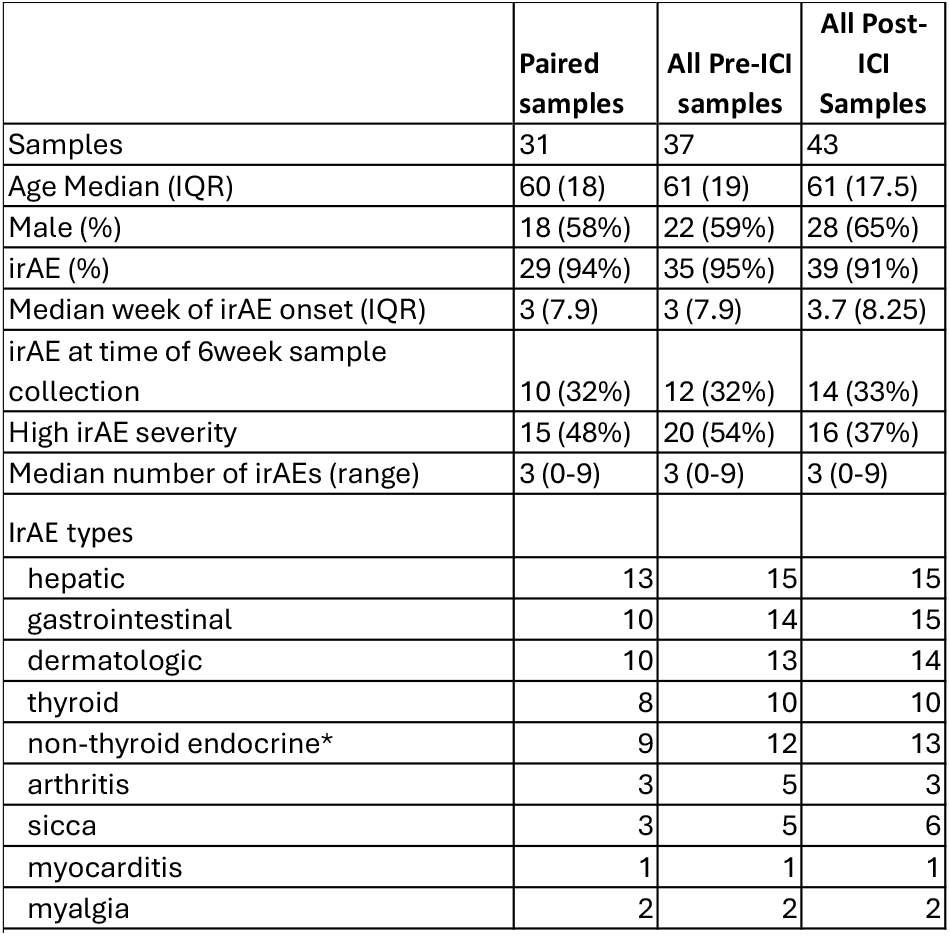
Description of all patients included in the mass cytrometry analysis. *adrenal insufficiency, hypophysitis, diabetes

After treatment initiation, participants were diagnosed with irAEs of varying severity, type, and onset time. The most common irAEs were gastrointestinal and endocrine related. The median number of irAEs per participant was 3. 61% of paired patients had already been diagnosed with at least one irAE at the 6-week PBMC collection timepoint, and the majority of participants (95%) were diagnosed with an irAE at some point during the trial. For analysis of baseline sample associations with irAE severity, we considered all irAEs diagnosed at any time following the initial dosing. irAEs were graded 0-5 using the Common Terminology Criteria for Adverse Events (CTCAE) and were divided into low (grade 1-2) and high (grade 3-5) severity (24).

We obtained PBMCs before treatment from 49 unique patients and paired blood 6 weeks after treatment initiation (after 2 doses of combination ICI therapy) from 31 of these patients. Cryopreserved PBMCs were processed for mass cytometry using a panel of antibodies to identify major lineages as well as specific T cell phenotypes (Figure 1A). Gated live cells, gated using FlowJo, were corrected for batch effects using the CyCombine R package (25) and then clustered in an unsupervised manner using Seurat (26). We then analyzed the data for effects of therapy and associations with irAE severity.

**Figure 1.**
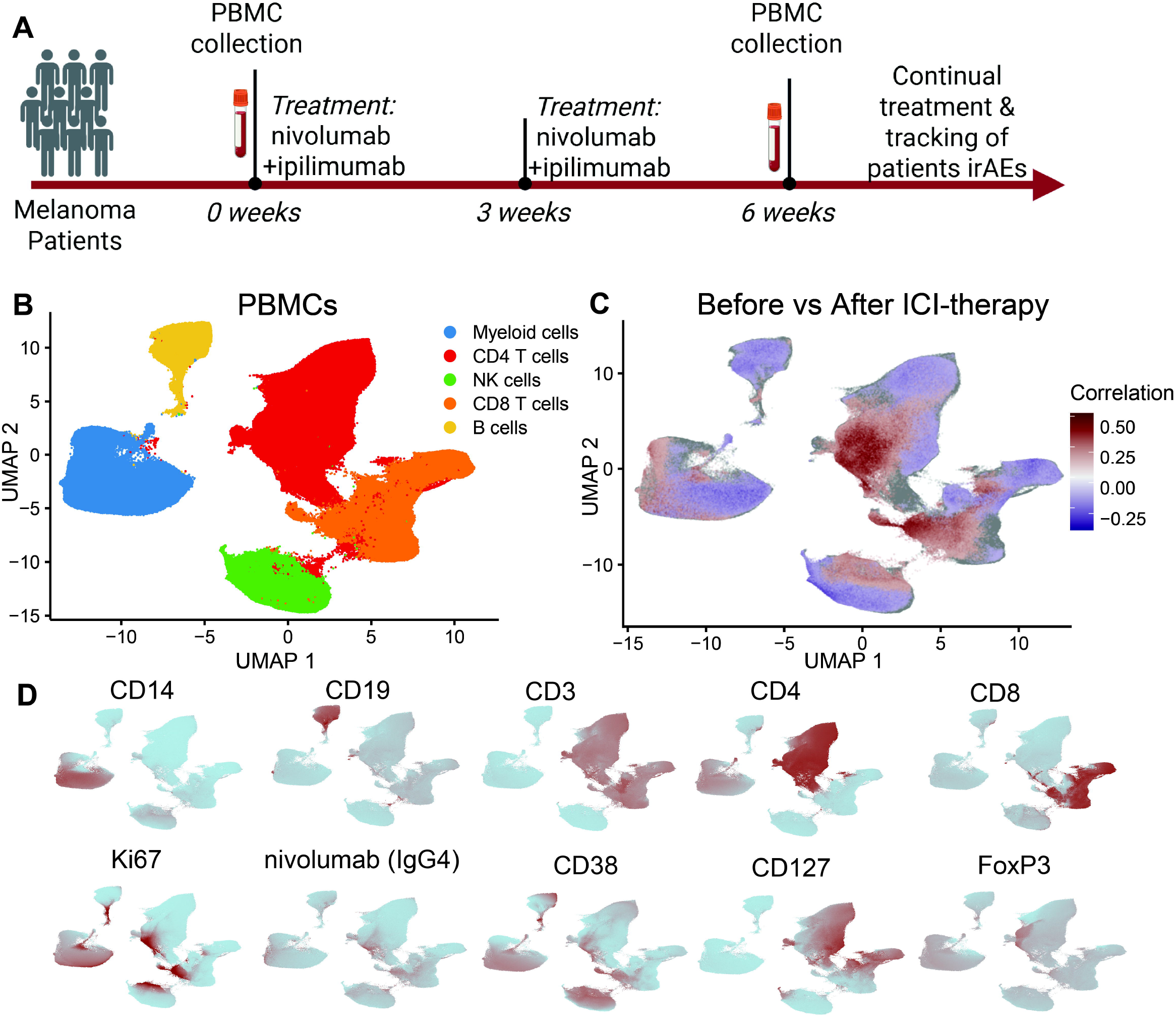
Changes in circulating cell frequency following combination ICI therapy. A) Schematic of experimental setup of analysis of PBMCs from patients with melanoma undergoing ICI treatment. B) UMAP representation following unsupervised clustering of PBMCs of melanoma patients before and after treatment (n = 49 patients; 31 paired). C) CNA analysis of before treatment samples vs after treatment samples. Red cells are significantly associated with post treatment, while blue cells are significantly associated with pre-treatment. Global p value = 0.023. D) Feature plots of the indicated markers on the PBMC UMAP. Red = high expression.

### Changes in proportions of immune cells following combination ICI therapy

We looked broadly at changes in frequencies of circulating immune cell populations induced by ICI therapy. The total proportions of CD4^+^ T cells, CD8^+^ T cells, NK cells, or myeloid cells did not consistently change following initiation of combination ICI therapy (Figure 1B, S1A-B). The frequency of total circulating B cells, however, was reduced following ICI therapy (Figure S1A). To gain further insight into specific B cell changes following checkpoint blockade and to determine any association with irAE severity in this cohort, we performed flow cytometry focused on B cell populations on the same samples. We confirmed a reduction of CD19^+^ cells in PBMCs in this dataset (Figure S2A). However, among CD19^+^ B cells, we observed increases in the proportion of both CD21lo B cells and CD38^hi^CD27^hi^ plasmablasts (Figure S2B-C). The frequency of CD27^+^ memory B cells did not change after therapy; however, IgG^+^ memory B cells decreased significantly in most patients (Figure S2D-E). Despite the changes in B cell frequencies, the baseline frequencies of each B cell population and the changes induced by ICI therapy in each B cell population were similar between those who were diagnosed with a severe irAE or a mild irAE (Figure S3A-D).

In addition to B cells, we also analyzed frequencies of plasmacytoid dendritic cells (pDCs) before and after combination ICI therapy by flow cytometry due to their association with autoimmune diseases and their production of type I interferon, a factor that promotes expansion of CD38^hi^ CD8^+^ T cells in ICI-associated arthritis (28). pDCs, identified as CD3^-^CD19^-^CD14^-^CD11C^-^ BDCA2^+^CD123^+^ cells, were decreased following ICI therapy, with similar decreases in patients with and without severe irAE development (Figure S2F, S2L). The frequency of pDCs at baseline was not associated with the irAE severity, and despite decreasing in most patients, the level of decrease in pDC frequency did not vary with irAE severity level (Figure S2M).

To assess effects of ICI therapy on circulating immune cell phenotypes broadly yet with high resolution, we utilized covarying neighborhood analysis (CNA) within the mass cytometry data (29). We observed significant differences in T cells, including clear increases in the post treatment group in subsets of CD4^+^ and CD8^+^ T cells (Figure 1C). Expanded T cells had phenotypes of memory T cell subsets characterized by high levels of CD38, low levels of CD127 and express the proliferation marker Ki67 (Figure 1D).

### Changes in circulating T cells following combination ICI therapy

To evaluate changes in T cells more granularly, we performed focused analyses on CD4^+^ T cells and CD8^+^ T cells (Figure 2A,2D and S3A-B). CNA and cluster analyses demonstrated widespread changes in both CD4^+^ and CD8^+^ T cell composition following ICI therapy (Figure 2B,2E). The most clearly expanded CD4^+^ T cells were in CD4 Cluster 10, which showed high levels of CD38, low levels of CD127, and also expression of granzyme B, granzyme K, perforin, and HLA-DR (Figure 2C, 2H, S3A). These CD4^+^ T cells were the highest expressors of the proliferation marker Ki67 (Figure 2C). Similarly, CD8 Cluster 6 which closely resembles CD4 Cluster 10 was dramatically induced following ICI therapy (Figure 2F, 2I), consistent with prior analyses (28). Interestingly, CD4 Cluster 8, which was characterized by granzyme B expression, but not CD38, did not change following ICI therapy (Figure 2B, 2C, S3C). Thus, cytotoxic CD4^+^ T cells in general did not increase, but rather a separate population of proliferating CD4s with granzyme B expression increased. We confirmed these changes to T cell populations using bi-axial gating (Figure S3E-F).

**Figure 2.**
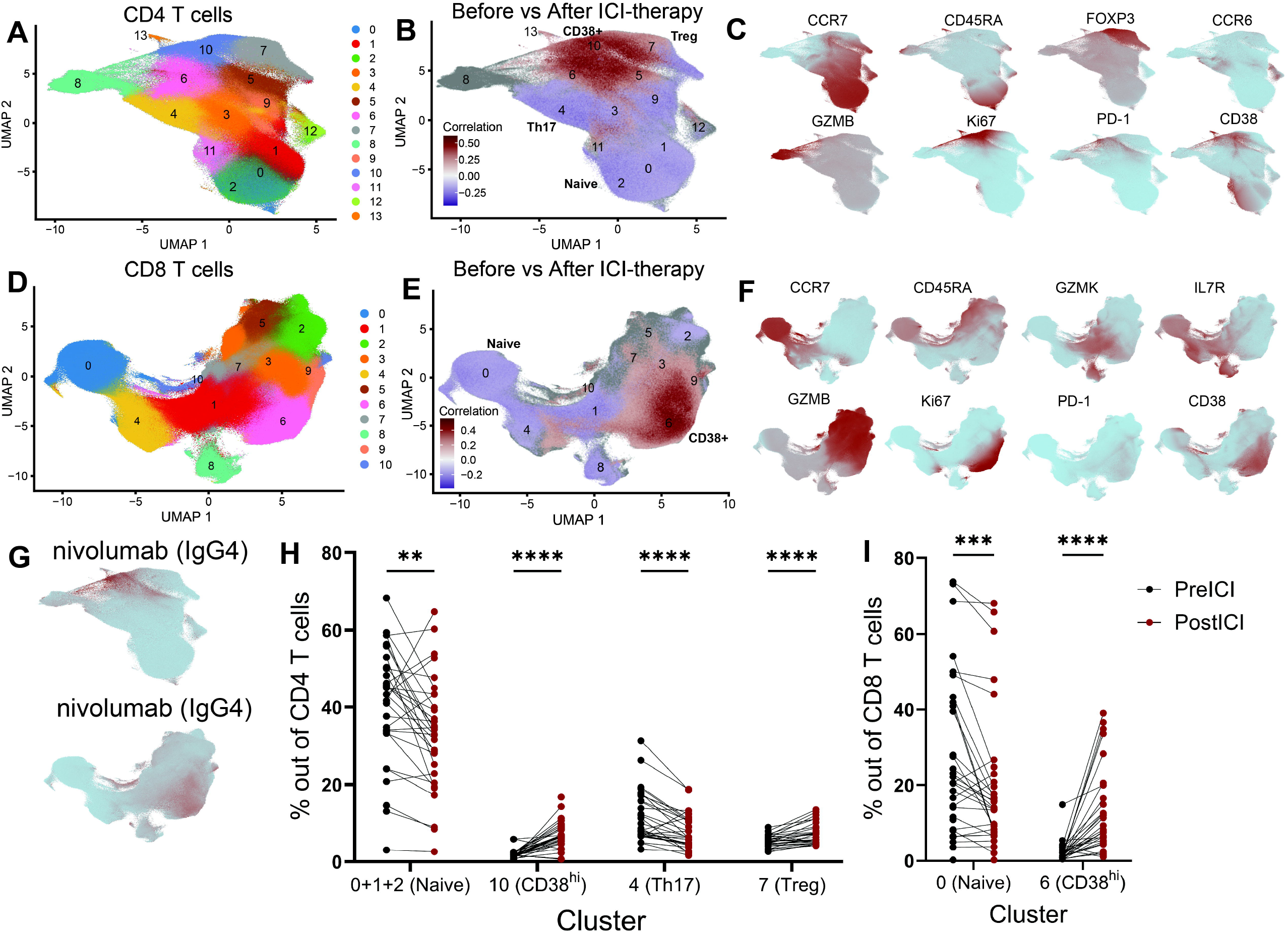
Changes in T cells following combination ICI therapy. A) UMAP representation following unsupervised clustering of CD4 T cells. B) CNA analysis of before treatment samples vs after treatment samples. Global p value = 0.001. C) Feature plots of the indicated markers on the CD4 T cell UMAP. D) UMAP representation following unsupervised clustering of CD8 T cells. E) CNA analysis of before treatment samples vs after treatment samples. Global p value = 0.001. F) Feature plots of the indicated markers on the CD8 T cell UMAP. G) Feature plots of in the CD4 (left) and CD8 (right) T cells indicating detection of nivolumab. H) Paired analysis of percent of indicated cluster in CD4 or I) CD8 T cells. **p<0.01, ***p<0.001, ****p<0.0001 by Wilcoxon paired test.

Our mass cytometry panel included an antibody against IgG4, the isotype of nivolumab (30), enabling detection and evaluation of cells bound by nivolumab. Most of the IgG4^+^ cells, the cells currently being treated by nivolumab, were identified in the expanded Ki67^hi^CD38^hi^ cluster. This observation was found in both the CD4 and CD8^+^ T cells (Figure 2G). Both CD8^+^ and CD4^+^ T cells were bound by nivolumab, although CD8^+^ T cells had a higher mean binding of nivolumab (Figure S6A-B). The frequency of nivolumab-bound CD4^+^ or CD8^+^ T cells was not higher in those patients with current or future severe irAEs (grade 3-4) compared to patients with less severe irAEs (grade 1-2) (Figure S6C). Similarly, the expression of PD-1 on T cells at baseline was not different between patients with irAEs of grades 0-2 irAEs (“Low Severity”) and grades 3-4 (irAEs (“High Severity”) and was not correlated with time of irAE onset or the total number of irAEs (Figure S6D). Other CD4^+^ T cell cluster changes included a decrease in the proportion of Cluster 4, characterized by CCR6 and CD161, representing a decrease in Th17 cells (Figure 2H, S3C-F). Additionally, both naïve CD4^+^ and CD8^+^ T cells exhibited a slight but significant decrease in proportion following ICI therapy (Figure 2H-I, S3C-F).

We also observed a significant increase in the frequency of Cluster 7 following combination ICI therapy. Cluster 7 was characterized by FOXP3, CD25, and Helios, consistent with CD4^+^ regulatory T cells (Tregs) (Figure 2H). In this cohort, ICI therapy induced an overall increase in frequency of CD25^hi^CD127^-^ CD4^+^ T cells (Figure S4A,C). Within Tregs, we observed clear changes in Treg phenotypes following ICI therapy (Figure S4B). Several markers related to Treg function and activation increased, including FOXP3, Helios, CD39, CD38, ICOS, HLA-DR, and Ki67, while TCF1 decreased (Figure S4D-E).

### Frequency of naïve T cells is associated with irAE severity

We next attempted to identify PBMC populations in the baseline, pre-treatment samples that were associated with severity of future irAE following ICI therapy. We first employed CNA to look for correlations of fine-grained neighborhoods with severity of subsequent irAEs (29). CNA analysis comparing PBMCs from patients who subsequently developed high grade irAEs (“High Severity”) to those who did not (“Low Severity”) identified several cell neighborhoods significantly correlated with irAE severity (Figure 3A). The cell neighborhoods with positive correlation values correlated with development of severe irAEs; these neighborhoods were primarily composed of naïve CD4^+^ and naïve CD8^+^ T cells (Figure 3A). Among CD4^+^ T cells, the association of naïve CD4^+^ T cells with development of severe irAEs remained even when adjusting for age and sex (Figure 3B, C). To confirm this association using a cell cluster-based analysis, we implemented mixed-effects modeling of associations of single cells (MASC), which identifies cell clusters associated with a clinical variable using a mixed-effect logistic regression model at a single-cell level. Consistent with CNA results, MASC analyses controlling for age and sex also identified CD4 Clusters 0 and 1 as significantly associated with severe irAEs, while CD4 Clusters 5, 6, 9, and 13 were negatively associated with severe irAEs (Figure 3D). CD4 Clusters 0 and 1 contained naïve CD4 T cells, while Clusters 5, 6, 9, and 13 contained memory CD4 populations including those expressing CD40L, Tbet, CD57, and granzyme B. Among CD8^+^ T cell clusters, MASC analysis highlighted the naïve CD8^+^ T cell cluster, cluster 0, as associated with development of severe irAEs; however, this observation did not pass correction for multiple testing (Figure S6E).

**Figure 3.**
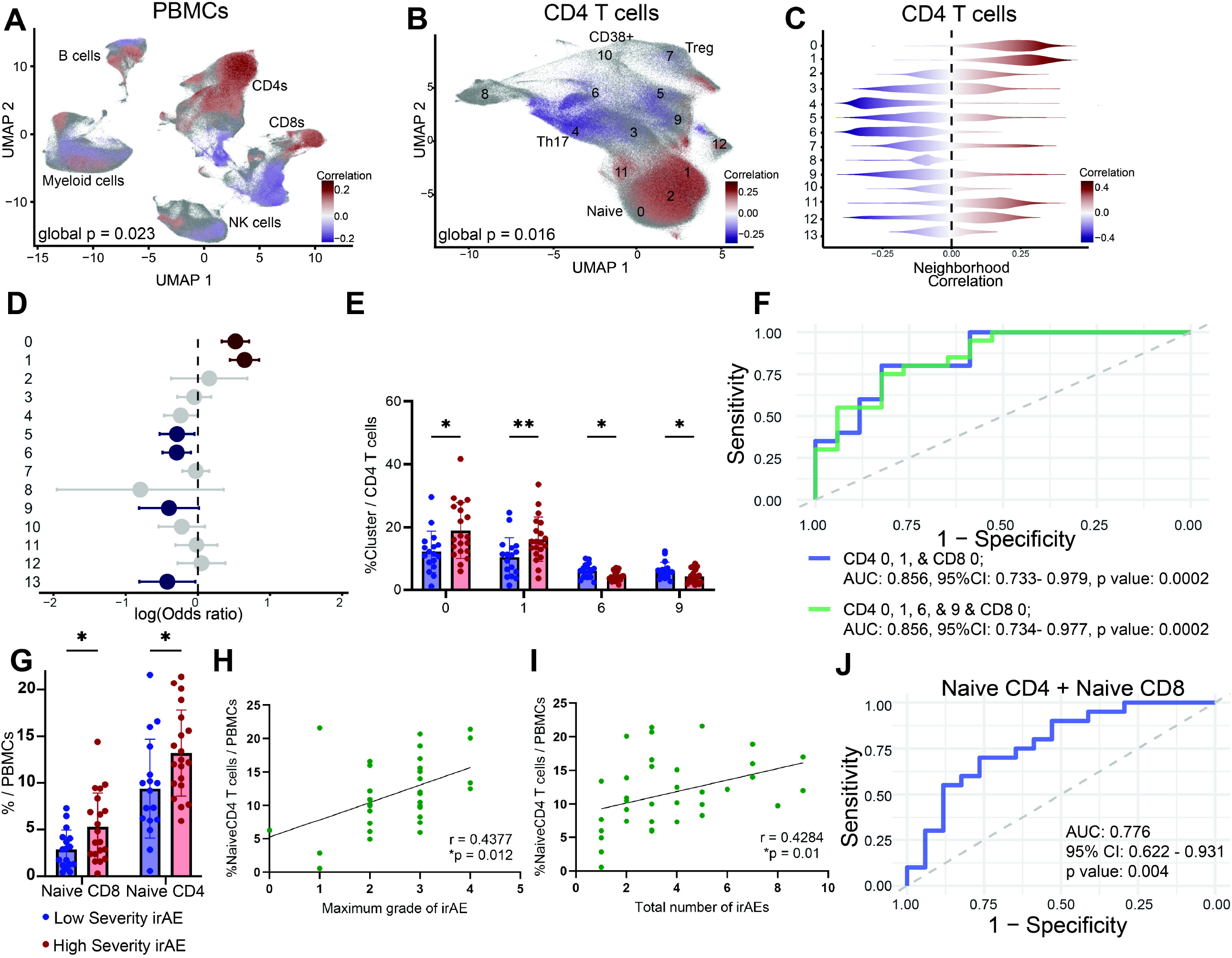
Naïve T cell levels before ICI therapy are associated with irAE severity. A) CNA analysis for high severity irAE PBMCs or B) CD4 T cells before ICI treatment. Red correlation values indicate significant positive association with development of severe irAE. All colored cells are filtered for FDR < 0.1. C) Correlation values of cells shown in B) sorted by cluster membership of the correlated cells. Red indicates positive correlation with severe irAEs. D) MASC analysis of CD4 T cells. Red indicates positive association with high severity irAE, and blue indicates association with low severity irAE. p<0.05 and are filtered for FDR < 0.1. Bars indicate confidence interval. E) Proportion of CD4 clusters at baseline with p < 0.05 for comparison of low vs high severity. F) ROC curves of proportion of CD4 clusters 0, 1, and CD8 cluster 0 (in blue) and CD4 clusters 0, 1, 6, and 9 and CD8 cluster 0 (in green) for development of severe irAE. G) Gating of mass cytometry data to quantify frequency of naïve T cells (CD3+CD4+CD45RA+CCR7- or CD3+CD8+CD45RA+CCR7-) out of total PBMCs of patients grouped by irAE severity. *p <0.05 by Mann Whitney test. H) Spearman correlation of gated naïve CD4 T cell proportion before ICI treatment with maximum grade or I) total number of irAEs (n= 37 patients). J) ROC curve of proportion of naïve CD4 T cells and naïve CD8 T cells for development of severe irAE.

The proportions of CD4^+^ cluster T cell clusters 0 and 1 out of all CD4 T cells were significantly higher at baseline in those who developed high severity irAEs, and the proportions of clusters 6 and 9 were significantly lower (Figure 3E). Additionally, the proportion of CD8 cluster 0 out of all CD8 T cells was significantly higher in those who developed high severity irAEs than those who developed low severity irAEs (Figure S6F). In ROC analysis, the AUC for the baseline proportion of CD4 T cell clusters 0, 1, 6, 9, and CD8 cluster 0 could predict severe irAE (AUC: 0.856, 95% CI: 0.734 - 0.934)(Figure 3F). The same analysis including naïve T cells only, CD4 T cell clusters 0, 1, and CD8 cluster 0, resulted in the same AUC (Figure 3F).

We next attempted to confirm these observations using bi-axial gating of the mass cytometry data. Bi-axial gating of the memory CD4 T cell subsets could not replicate the clusters associated with lower severity irAEs (clusters 6,9); therefore we did not pursue these further. However, naïve CD4^+^ or CD8^+^ T cells, corresponding to CD4 clusters 0 and 1 and CD8 cluster 0, could be easily gated as CD19^-^CD14^-^CD3^+^ CD4^+^/CD8^+^ CD45RA^+^ CCR7^+^. Patients who developed severe irAEs had a higher proportion of gated naïve CD4 and CD8^+^ T cells out of all PBMCs at baseline than patients who did not develop severe irAEs (Figure 3G). Further, the frequency of all naïve CD4^+^ T cells (r = 0.4377, p = 0.012) and naïve CD8^+^ T cells (r = 0.3854, p = 0.029) before therapy was significantly correlated with maximum grade of future irAE (Figure 3H, S6C). Frequency of naïve CD4^+^ T cells, but not naïve CD8^+^ T cells, at baseline was also correlated with the total number of irAEs per patient (r = 0.4284, p = 0.01) (Figure 3I, Figure S6D). ROC analysis of gated naïve CD4 and CD8^+^ T cells predicted onset of severe irAE (AUC: 0.776, 95% CI: 0.622 – 0.931) (Figure 3J). Naïve CD4 T cell proportion at baseline was not associated with overall survival of melanoma within this cohort (Figure S6F-G)(31).

## Discussion

In this study, we utilized high-dimensional mass cytometry analyses to assess both changes in circulating immune cell profiles induced by combination PD-1/CTLA-4 blockade and cellular features associated with the development of severe irAEs in treated patients with melanoma. Understanding the features of systemic immune activation and function altered by checkpoint blockade, as well as the specific immune features associated with development of irAE, is critical in developing a more precise prediction of potential effects and consequences of checkpoint inhibitor therapies.

We demonstrated that the frequency of naïve T cells, especially naïve CD4^+^ T cells, before treatment is associated with the development of a severe irAE in this cohort. These results differ somewhat from those in an observational cohort study of patients with melanoma treated with either PD-1/CTLA-4 blockade or PD-1 blockade reported by Lozano et al (22), which indicated an association of an increased proportion of effector memory CD4^+^ T cells among total PBMC with development of severe irAEs. However, in the study by Lozano et al, multiple CD4 T cell populations, including naïve CD4^+^ T cells, quantified among total PBMC, showed the same direction of effect as did naïve CD4^+^ T cells in our study. Our study refines the specific CD4^+^ T cell population most associated with severe irAEs, here using analysis of a cohort of patients treated with a uniform ICI therapy in the context of a clinic trial. Although we did not have TCR sequencing in our study, the observation that increased TCR repertoire diversity is also associated with more severe irAEs (22) is conceptually consistent with the findings in our study. An increased number of naïve T cells should correspond with an increased overall TCR repertoire diversity and represents a younger or ‘less experienced’ immune system. This observation is also consistent with a separate prior observational study of 28 patients with melanoma treated with different ICI therapies that also indicated an association of naïve T cells at baseline with development of severe irAEs, as well as clinical benefit (21).

Profiling T cells before and after combination PD-1/CTLA-4 therapy demonstrated a large set of prominent changes in circulating T cell phenotypes, including marked expansion of activated, CD38^hi^ Ki67^+^ T cells among both CD4^+^ T cells and CD8^+^ T cells. The inclusion of an anti-IgG4 antibody in the staining panel enabled detection and evaluation of cells directly bound by nivolumab. These nivolumab-bound CD4^+^ T cells are highly enriched in activated and cytotoxic CD4^+^ T cell phenotypes, consistent with our prior analyses (28), suggesting that T cells released from PD-1/CTLA-4 inhibition acquire an activated, proliferating phenotype systemically. A recent study corroborates our findings of an expansion of Ki67^+^CD38^hi^ T cells early following ICI therapy and identifies these cells as expanded early after treatment in those with irAEs (32). Importantly, our data does not show a link between frequency of nivolumab-bound T cells and irAE severity.

While some studies indicate that Tregs are depleted in the tumor microenvironment following CTLA-4 blockade, their change in frequency in circulation has been less well characterized, especially following therapy of both anti-CTLA-4 and anti-PD-1 (33). We identified an increase in FOXP3+ Tregs following combination PD-1/CTLA-4 blockade, with the expanded Tregs showing increased expression of Treg and activation markers. These changes in Tregs may be due to CTLA-4 blockade, PD-1 blockade, or both. While we did not include an antibody to detect cell-bound ipilimumab in our staining panel, it is plausible that CTLA-4 blockade affects Treg numbers and phenotype. Ipilimumab can induce antibody-dependent cellular cytotoxicity and phagocytosis on intratumoral Tregs (34-38); however, the absolute numbers of intratumoral Tregs increases in melanoma tumors after CTLA-4 blockade (39). Absence of PD-1 on Tregs can also change Treg features and function, including through upregulation of CD30 expression (40). Future work should include gene expression level analysis on Tregs before and after ICI therapy to further characterize changes to individual Tregs populations, such as CD25^hi^ or CD25^low^ Tregs.

This cohort is made up of patients with a single malignancy treated with one specific protocol of combination PD-1/CTLA-4 blockade. While our results cannot be extrapolated to other treatment protocols at this time, our analysis is strengthened by lack of extraneous heterogeneity. Our study is limited by one cohort of patients and thus should be validated within separate cohorts and methods in the future. Additionally, in this cohort, we cannot assess cellular features associated with a lack of irAE development because almost all of the patients studied developed at least 1 mild irAE. The study was not powered to evaluate organ-specific irAE manifestations, and the multiple irAEs that occur in individual patients complicate testing for cellular associations with individual organ-specific irAE. The cytometric profiling approach focused primarily on lymphocyte features; thus, we have not fully captured the range of changes on myeloid and dendritic cell populations. Nonetheless, the use of high dimensional profiling of longitudinal samples from a clinical trial provide a robust, broad assessment of cellular phenotypes associated with checkpoint blockade in this setting. Our study specifically associates circulating naïve CD4^+^ T cells with predisposition to high severity irAE.

## Methods

### Study approval and design

Human subjects research was performed in accordance with the Institutional Review Board at Memorial Sloan Kettering and Mass General Brigham. Blood samples were obtained through the ADAPT-IT trial from patients with melanoma before and after receiving 2 doses of both nivolumab and ipilimumab (5). Demographics of each patient can be found in Supplementary Table 1. Mononuclear cells from PBMCs were isolated by density centrifugation using Ficoll-Paque Plus (GE healthcare) and cryopreserved in 10% DMSO in FBS and stored in liquid nitrogen.

### Sex as a biological variable

We included both male and female patient samples in our analysis. We have included sex as a covariate in our statistical analyses since proportions of naïve T cells especially CD8^+^ T cells can be associated with age (Figures S7A-B). We have noted this in the corresponding figure legends.

### Mass cytometry of ICI-treated PBMCs

Cryopreserved PBMCs were thawed, washed and counted for mass cytometry staining. 2 million live PBMCs were stained for mass cytometry from each sample. A total of 86 samples were stained in 5 batches. Additionally, a reference sample isolated from a leukoreduction collar was included in each batch. Antibodies were acquired from Fluidigm and Longwood Medical Area CyTOF Core. Clones of the antibodies can be found in Supplementary Table 2. Cells were stained for viability using 103Rh, followed by FC block (Invitrogen). Surface staining was done in cell staining buffer (Fluidigm) for 30 minutes followed by fixing and permeabilization with FoxP3/Transcription Factor Staining Buffer Set (Invitrogen) and barcoding (Fluidigm). Intracellular staining was done on pooled barcoded samples. Following fixing in 4% PFA, cells were diluted in CyTOF PBS containing appropriate concentration of Intercalator-Ir, washed and resuspended in cell acquisition plus buffer containing EQ beads (Fluidigm). Cells were acquired on a CyTOF-XT (Fluidigm).

### Mass cytometry data analysis of ICI-treated PBMCs

Cytometry data was normalized and debarcoded as previously described (41). Live cells were gated using FlowJo (BD) as DNA+103Rh-Beads-events. Batch correction on total live cells was performed by the cyCombine R package (25). Following batch correction, 15,000 cells per sample were analyzed in an unsupervised manner using the Seurat pipeline for R. Further characterization of CD8^+^ and CD4^+^ T cell populations was carried out by gating for CD3+CD8+CD4- and CD3+CD8-CD4+ cells respectively followed by unsupervised analysis using the Seurat R package. CD4^+^ and CD8^+^ cluster resolutions were chosen based on ability to identify clear definitions of CD4^+^ or CD8^+^ T cell subsets and then confirmed using the ClusTree package. Analysis by biaxial gating was done on live batch corrected cells using FlowJo (BD).

### Statistical analysis of ICI-treated PBMCs

Comparisons of frequencies of clusters of cells before and after therapy was done using Wilcoxon paired test in GraphPad Prism. irAE severity was considered high if greater than or equal to a max grade of 3. Comparisons of frequencies of cells between high severity and low severity irAE patients was done with a Mann Whitney test in GraphPad Prism.

To identify cell clusters associated with high severity irAEs at the single cell level we utilized mixed-effects association of single cells (MASC) as previously described (42). To identify cells belonging to neighborhoods associated with high severity irAEs, we utilized Covarying Neighborhood Analysis (CNA) also as previously described (29). In both types of analysis cells or clusters were considered significantly associated if they had an adjusted p value of less than 0.1. ROC analysis was completed using the R package pROC and function glm. Analysis was completed for T cell cluster proportions and T cell proportions identified by bi-axial gating.

### Flow cytometry staining and analysis of ICI-treated PBMCs

Cryopreserved PBMCs were thawed, counted and washed as before. Approximately 1 million PBMCs were stained for each of 86 samples in 5 batches. Antibodies included anti-CD14 AF350, anti-CD19 APC-Cy7, anti-CD20 BV605, anti-CD27 BV421, anti-CD38 BV786, anti-CD21 APC, anti-PE CD11C, anti-IgD Percp-cy5.5, anti-IgG BV510, anti-IgA FITC, anti-IgM PE-Cy7, CD24 BV711, anti-BDCA2 BUV737 (BD Biosciences), anti-CD123 BV650 (all Biolegend unless otherwise indicated). Analysis by biaxial gating was performed using FlowJo (BD).

## Data availability

Raw data will be uploaded as required by journal at location which will be written here following acceptance. R codes for analysis will be made available upon request.

## Author contributions

DAR, AB, LD, and KEM conceptualized the project. KEM, ME, and IA performed experiments. KEM analyzed data. AH and NG advised data analysis. MP provided patient samples. KEM and DAR drafted the manuscript, and all authors edited the manuscript.

## Acknowledgements

Funding for this study was provided in part by an Innovative Research Award from the Rheumatology Research Foundation (to DAR, AB, LD), a Career Award in Medical Sciences from the Burroughs Wellcome Fund (to DAR), and NIAMS P30 AR070253. M.A.P. would like to acknowledge institutional support from the National Cancer Institute/ National Institutes of Health Cancer Center Support Grant (P30CA008748).

## Figure Legends

**Supplementary Figure 1**. A) Paired analysis of frequency of major cell clusters before vs after treatment per patient. * p<0.05 by Wilcoxon paired test. B) DotPlot of all mass cytometry markers in PBMCs by low resolution clusters (see Figure 1B-D).

**Supplementary Figure 2**. A - E) Paired analysis of frequency of indicated B cell markers analyzed by flow cytometry before vs after treatment per patient in indicated populations. F) Paired analysis of frequency of plasmacytoid dendritic cells (gated as CD3-CD19-CD14-CD11C-BDCA2+CD123+) before vs after treatment per patient. A G, H) Analysis of frequency of indicated B cell populations analyzed by flow cytometry before treatment per patient in indicated populations. Frequencies of B cells in patients with low and high severity irAEs are compared. I-K) Change in frequency of B cell populations after - before treatment. L) Frequency of plasmacytoid dendritic cells (gated as CD3-CD19-CD14-CD11C-BDCA2+CD123+). M) Change in frequency of plasmacytoid dendritic cells after - before treatment. Red bars indicate patients with high severity irAEs, and blue bars indicate patients with low severity irAEs. -F: * p<0.05, ** p < 0.01, *** p <0.001 by Wilcoxon paired test. G-M: All plots are not significant by Mann Whitney test.

**Supplementary Figure 3**. A) DotPlot representation of all mass cytometry markers in CD4+ T cells subset from PBMCs (see Figure 2A-C). B) Dot Plot representation of all mass cytometry markers in CD8+ T cells subset from PBMCs (see Figure 2D-F). C) Paired analysis of frequency of all CD4 T cell and D) CD8 T cell clusters before vs after treatment per patient. test. E) Paired analysis of CD4 and F) CD8 T cells analyzed by biaxial gating. * p<0.05, **p<0.01, ***p<0.001, ****p<0.0001 by Wilcoxon paired analysis.

**Supplementary Figure 4**. A) Paired analysis of frequency of CD25^hi^CD127^-^ CD4 T cells before and after ICI therapy by biaxial gating. B) UMAP representation following unsupervised clustering of CD4 regulatory T cells. C) CNA analysis of before treatment samples vs after treatment samples. Global p value = 0.001. D) Feature plots of the indicated markers on the Treg UMAP. E) Paired analysis of Median Fluorescent Intensity of FoxP3 and Helios or F) Ki67, ICOS, CD39, HLA-DR, and CD38 in CD25^hi^CD127^-^ CD4 T cells before and after ICI therapy. ***p<0.001, ****p<0.0001 by Wilcoxon paired analysis.

**Supplementary Figure 5**. A) Frequency of IgG4^+^PD-1^+^ CD4^+^ or CD8^+^ T cells out of CD3^+^ T cells by gating of mass cytometry analysis in the post-treatment samples. B) Frequency of IgG4^+^PD-1^+^ out of CD4^+^ or CD8^+^ T cells by gating of mass cytometry analysis in the post-treatment samples. C) Frequency of IgG4^+^PD-1^+^ out of CD4^+^ (right) or CD8^+^ (left) T cells in patients with low or high severity of irAE by gating of mass cytometry analysis. D) Frequency of total PD-1^+^ out of CD4^+^ (right) or CD8^+^ (left) T cells in pre-treatment samples from patients with low or high severity of irAE by gating of mass cytometry analysis.

**Supplementary Figure 6**. A) Spearman correlation of proportion of gated naïve CD4+ T cells (CD19^-^CD14^-^ CD3^+^CD4^+^CD45RA^+^CCR7^+^) with age. B) Spearman correlation of proportion of gated naïve CD8+ T cells (CD19^-^CD14^-^ CD3^+^CD8^+^CD45RA^+^CCR7^+^) with age, C) maximum grade or irAE, and D) total number of irAEs diagnosed. E) MASC analysis of CD8 T cell clusters (see Figure 2). + indicates p < .05, adj p = 0.5. F) Proportion of cluster 0 out of CD8 T cells. p<0.05 by Mann Whitney test. G) MASC analysis for overall survival in CD4^+^ and H) CD8^+^ T cells before treatment (survived n = 5, not survived n = 29).

## References

1. Havel JJ, Chowell D, and Chan TA. The evolving landscape of biomarkers for checkpoint inhibitor immunotherapy. Nat Rev Cancer. 2019;19(3):133–50.

2. Schadendorf D, Hodi FS, Robert C, Weber JS, Margolin K, Hamid O, et al. Pooled Analysis of Long-Term Survival Data From Phase II and Phase III Trials of Ipilimumab in Unresectable or Metastatic Melanoma. J Clin Oncol. 2015;33(17):1889–94.

3. Hodi FS, O’Day SJ, McDermott DF, Weber RW, Sosman JA, Haanen JB, et al. Improved survival with ipilimumab in patients with metastatic melanoma. N Engl J Med. 2010;363(8):711–23.

4. Ribas A, and Wolchok JD. Cancer immunotherapy using checkpoint blockade. Science. 2018;359(6382):1350–5.

5. Postow MA, Goldman DA, Shoushtari AN, Betof Warner A, Callahan MK, Momtaz P, et al. Adaptive Dosing of Nivolumab + Ipilimumab Immunotherapy Based Upon Early, Interim Radiographic Assessment in Advanced Melanoma (The ADAPT-IT Study). J Clin Oncol. 2022;40(10):1059–67.

6. Wei SC, Anang N-AAS, Sharma R, Andrews MC, Reuben A, Levine JH, et al. Combination anti–CTLA-4 plus anti– PD-1 checkpoint blockade utilizes cellular mechanisms partially distinct from monotherapies. Proceedings of the National Academy of Sciences. 2019;116(45):22699–709.

7. Das R, Verma R, Sznol M, Boddupalli CS, Gettinger SN, Kluger H, et al. Combination therapy with anti-CTLA-4 and anti-PD-1 leads to distinct immunologic changes in vivo. J Immunol. 2015;194(3):950–9.

8. Das R, Bar N, Ferreira M, Newman AM, Zhang L, Bailur JK, et al. Early B cell changes predict autoimmunity following combination immune checkpoint blockade. The Journal of Clinical Investigation. 2018;128(2):715–20.

9. Patrinely JR, Jr., Johnson R, Lawless AR, Bhave P, Sawyers A, Dimitrova M, et al. Chronic Immune-Related Adverse Events Following Adjuvant Anti-PD-1 Therapy for High-risk Resected Melanoma. JAMA Oncol. 2021;7(5):744–8.

10. Cappelli LC, and Bingham CO. Spectrum and impact of checkpoint inhibitor-induced irAEs. Nature Reviews Rheumatology. 2021;17(2):69–70.

11. Postow MA, Chesney J, Pavlick AC, Robert C, Grossmann K, McDermott D, et al. Nivolumab and Ipilimumab versus Ipilimumab in Untreated Melanoma. New England Journal of Medicine. 2015;372(21):2006–17.

12. Xu C, Chen Y-P, Du X-J, Liu J-Q, Huang C-L, Chen L, et al. Comparative safety of immune checkpoint inhibitors in cancer: systematic review and network meta-analysis. BMJ. 2018;363:k4226.

13. Martins F, Sofiya L, Sykiotis GP, Lamine F, Maillard M, Fraga M, et al. Adverse effects of immune-checkpoint inhibitors: epidemiology, management and surveillance. Nature Reviews Clinical Oncology. 2019;16(9):563–80.

14. Chennamadhavuni A, Abushahin L, Jin N, Presley CJ, and Manne A. Risk Factors and Biomarkers for Immune-Related Adverse Events: A Practical Guide to Identifying High-Risk Patients and Rechallenging Immune Checkpoint Inhibitors. Front Immunol. 2022;13:779691.

15. Suijkerbuijk KPM, van Eijs MJM, van Wijk F, and Eggermont AMM. Clinical and translational attributes of immune-related adverse events. Nature Cancer. 2024.

16. Kartolo A, Sattar J, Sahai V, Baetz T, and Lakoff JM. Predictors of Immunotherapy-Induced Immune-Related Adverse Events. Current Oncology. 2018;25(5):403–10.

17. Wang J, Ma Y, Lin H, Wang J, and Cao B. Predictive biomarkers for immune-related adverse events in cancer patients treated with immune-checkpoint inhibitors. BMC Immunology. 2024;25(1):8.

18. Shah KP, Song H, Ye F, Moslehi JJ, Balko JM, Salem J-E, et al. Demographic Factors Associated with Toxicity in Patients Treated with Anti–Programmed Cell Death-1 Therapy. Cancer Immunology Research. 2020;8(7):851–5.

19. Wong SK, Blum SM, Sun X, Da Silva IP, Zubiri L, Ye F, et al. Efficacy and safety of immune checkpoint inhibitors in young adults with metastatic melanoma. European Journal of Cancer. 2023;181:188–97.

20. van Eijs MJM, Verheijden RJ, van der Wees SA, Nierkens S, van Lindert ASR, Suijkerbuijk KPM, et al. Toxicity-specific peripheral blood T and B cell dynamics in anti-PD-1 and combined immune checkpoint inhibition. Cancer Immunology, Immunotherapy. 2023;72(12):4049–64.

21. Kovacsovics-Bankowski M, Sweere JM, Healy CP, Sigal N, Cheng L-C, Chronister WD, et al. Lower frequencies of circulating suppressive regulatory T cells and higher frequencies of CD4^+^ naïve T cells at baseline are associated with severe immune-related adverse events in immune checkpoint inhibitor-treated melanoma. Journal for ImmunoTherapy of Cancer. 2024;12(1):e008056.

22. Lozano AX, Chaudhuri AA, Nene A, Bacchiocchi A, Earland N, Vesely MD, et al. T cell characteristics associated with toxicity to immune checkpoint blockade in patients with melanoma. Nat Med. 2022;28(2):353–62.

23. Smithy JW, Kalvin HL, Ehrich FD, Shah R, Adamow M, Raber V, et al. Early On-Treatment Assessment of T Cells, Cytokines, and Tumor DNA with Adaptively Dosed Nivolumab + Ipilimumab: Final Results from the Phase 2 ADAPT-IT Study. Clinical Cancer Research. 2024;30(16):3407–15.

24. Minasian LM, O’Mara A, and Mitchell SA. Clinician and Patient Reporting of Symptomatic Adverse Events in Cancer Clinical Trials: Using CTCAE and PRO-CTCAE(®) to Provide Two Distinct and Complementary Perspectives. Patient Relat Outcome Meas. 2022;13:249–58.

25. Pedersen CB, Dam SH, Barnkob MB, Leipold MD, Purroy N, Rassenti LZ, et al. cyCombine allows for robust integration of single-cell cytometry datasets within and across technologies. Nature Communications. 2022;13(1):1698.

26. Hao Y, Hao S, Andersen-Nissen E, Mauck WM, Zheng S, Butler A, et al. Integrated analysis of multimodal single-cell data. Cell. 2021;184(13):3573-3573.e29.

27. Barth DA, Stanzer S, Spiegelberg JA, Bauernhofer T, Absenger G, Szkandera J, et al. Patterns of Peripheral Blood B-Cell Subtypes Are Associated With Treatment Response in Patients Treated With Immune Checkpoint Inhibitors: A Prospective Longitudinal Pan-Cancer Study. Frontiers in Immunology. 2022;13.

28. Wang R, Singaraju A, Marks KE, Shakib L, Dunlap G, Adejoorin I, et al. Clonally expanded CD38^hi^ cytotoxic CD8 T cells define the T cell infiltrate in checkpoint inhibitor–associated arthritis. Science Immunology. 2023;8(85):eadd1591.

29. Reshef YA, Rumker L, Kang JB, Nathan A, Korsunsky I, Asgari S, et al. Co-varying neighborhood analysis identifies cell populations associated with phenotypes of interest from single-cell transcriptomics. Nature Biotechnology. 2022;40(3):355–63.

30. Osa A, Uenami T, Koyama S, Fujimoto K, Okuzaki D, Takimoto T, et al. Clinical implications of monitoring nivolumab immunokinetics in non-small cell lung cancer patients. JCI Insight. 2018;3(19).

31. Cook S, Samuel V, Meyers DE, Stukalin I, Litt I, Sangha R, et al. Immune-Related Adverse Events and Survival Among Patients With Metastatic NSCLC Treated With Immune Checkpoint Inhibitors. JAMA Netw Open. 2024;7(1):e2352302.

32. Nuñez NG, Berner F, Friebel E, Unger S, Wyss N, Gomez JM, et al. Immune signatures predict development of autoimmune toxicity in patients with cancer treated with immune checkpoint inhibitors. Med. 2023;4(2):113-29.e7.

33. Wei SC, Levine JH, Cogdill AP, Zhao Y, Anang NAS, Andrews MC, et al. Distinct Cellular Mechanisms Underlie Anti-CTLA-4 and Anti-PD-1 Checkpoint Blockade. Cell. 2017;170(6):1120–33 e17.

34. Kavanagh B, O’Brien S, Lee D, Hou Y, Weinberg V, Rini B, et al. CTLA4 blockade expands FoxP3+ regulatory and activated effector CD4+ T cells in a dose-dependent fashion. Blood. 2008;112(4):1175–83.

35. Hong MMY, and Maleki Vareki S. Addressing the Elephant in the Immunotherapy Room: Effector T-Cell Priming versus Depletion of Regulatory T-Cells by Anti-CTLA-4 Therapy. Cancers (Basel). 2022;14(6).

36. Rosskopf S, Leitner J, Zlabinger GJ, and Steinberger P. CTLA-4 antibody ipilimumab negatively affects CD4+ T-cell responses in vitro. Cancer Immunology, Immunotherapy. 2019;68(8):1359–68.

37. Romano E, Kusio-Kobialka M, Foukas PG, Baumgaertner P, Meyer C, Ballabeni P, et al. Ipilimumab-dependent cellmediated cytotoxicity of regulatory T cells ex vivo by nonclassical monocytes in melanoma patients. Proceedings of the National Academy of Sciences. 2015;112(19):6140–5.

38. Simpson TR, Li F, Montalvo-Ortiz W, Sepulveda MA, Bergerhoff K, Arce F, et al. Fc-dependent depletion of tumorinfiltrating regulatory T cells co-defines the efficacy of anti–CTLA-4 therapy against melanoma. Journal of Experimental Medicine. 2013;210(9):1695–710.

39. Sharma A, Subudhi SK, Blando J, Scutti J, Vence L, Wargo J, et al. Anti-CTLA-4 Immunotherapy Does Not Deplete FOXP3+ Regulatory T Cells (Tregs) in Human Cancers. Clinical Cancer Research. 2019;25(4):1233–8.

40. Lim JX, McTaggart T, Jung SK, Smith KJ, Hulme G, Laba S, et al. PD-1 receptor deficiency enhances CD30+ Treg cell function in melanoma. Nature immunology. 2025;26(7):1074–86.

41. Bocharnikov AV, Keegan J, Wacleche VS, Cao Y, Fonseka CY, Wang G, et al. PD-1hi CXCR5-T peripheral helper cells promote B cells responses in lupus via MAF and IL-21. JCI insight. 2019.

42. Fonseka CY, Rao DA, Teslovich NC, Korsunsky I, Hannes SK, Slowikowski K, et al. Mixed-effects association of single cells identifies an expanded effector CD4(+) T cell subset in rheumatoid arthritis. Science translational medicine. 2018;10(463).

