## SupplementaryFigures for "Pre-treatment naïve T cells are associated with severe irAE following PD-1/CTLA4 checkpoint blockade for melanoma"

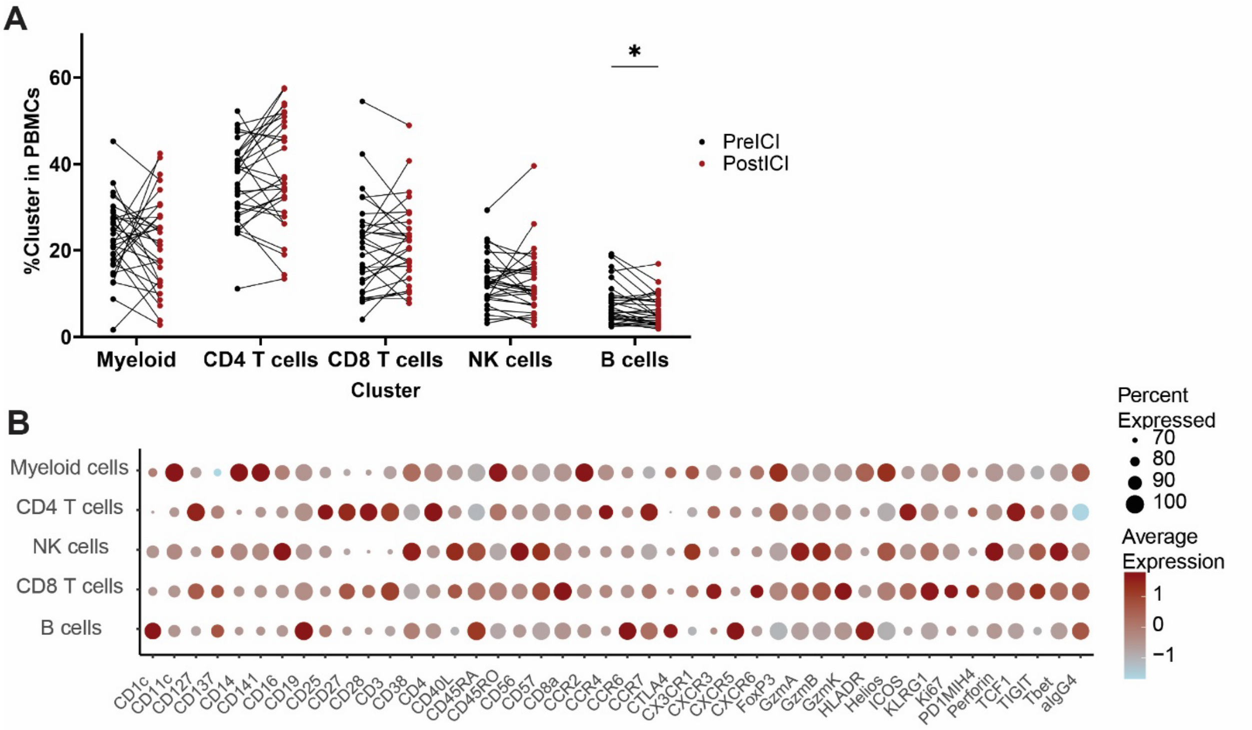


**Supplementary Figure 1**

A) Paired analysis of frequency of major cell clusters before vs after treatment per patient. * p<0.05 by Wilcoxon paired test. B) DotPlot of all mass cytometry markers in PBMCs by low resolution clusters (see Figure 1B-D).

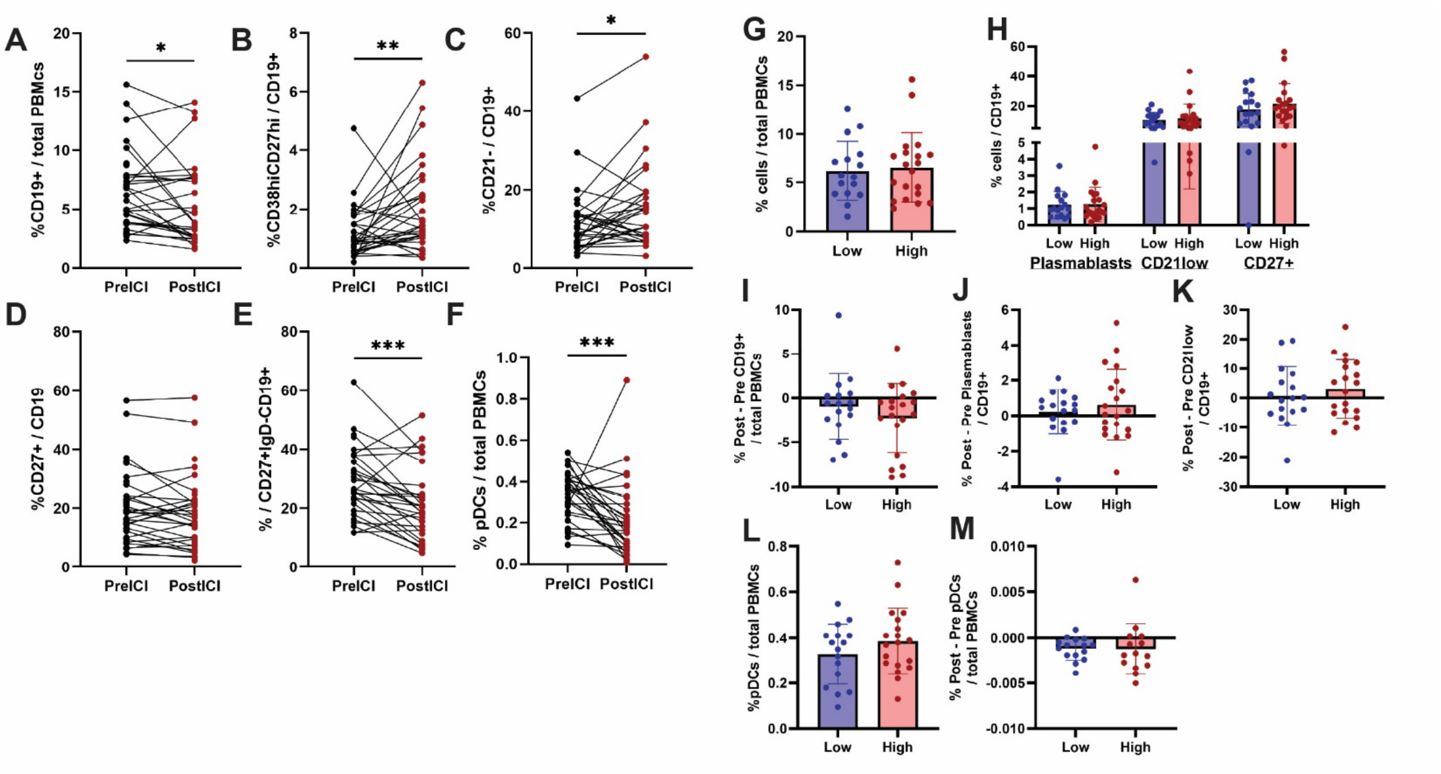

**Supplementary Figure 2. Changes to B cell frequencies following ICI treatment.** A - E) Paired analysis of frequency of indicated B cell markers analyzed by flow cytometry before vs after treatment per patient in indicated populations. F) Paired analysis of frequency of plasmacytoid dendritic cells (gated as CD3-CD19-CD14-CD11C-BDCA2+CD123+) before vs after treatment per patient. A G, H) Analysis of frequency of indicated B cell populations analyzed by flow cytometry before treatment per patient in indicated populations. Frequencies of B cells in patients with low and high severity irAEs are compared. I-K) Change in frequency of B cell populations after - before treatment. L) Frequency of plasmacytoid dendritic cells (gated as CD3-CD19-CD14-CD11C-BDCA2+CD123+). M) Change in frequency of plasmacytoid dendritic cells after - before treatment. Red bars indicate patients with high severity irAEs, and blue bars indicate patients with low severity irAEs. -F: * p<0.05, ** p < 0.01, *** p <0.001 by Wilcoxon paired test. G-M: All plots are not significant by Mann Whitney test.

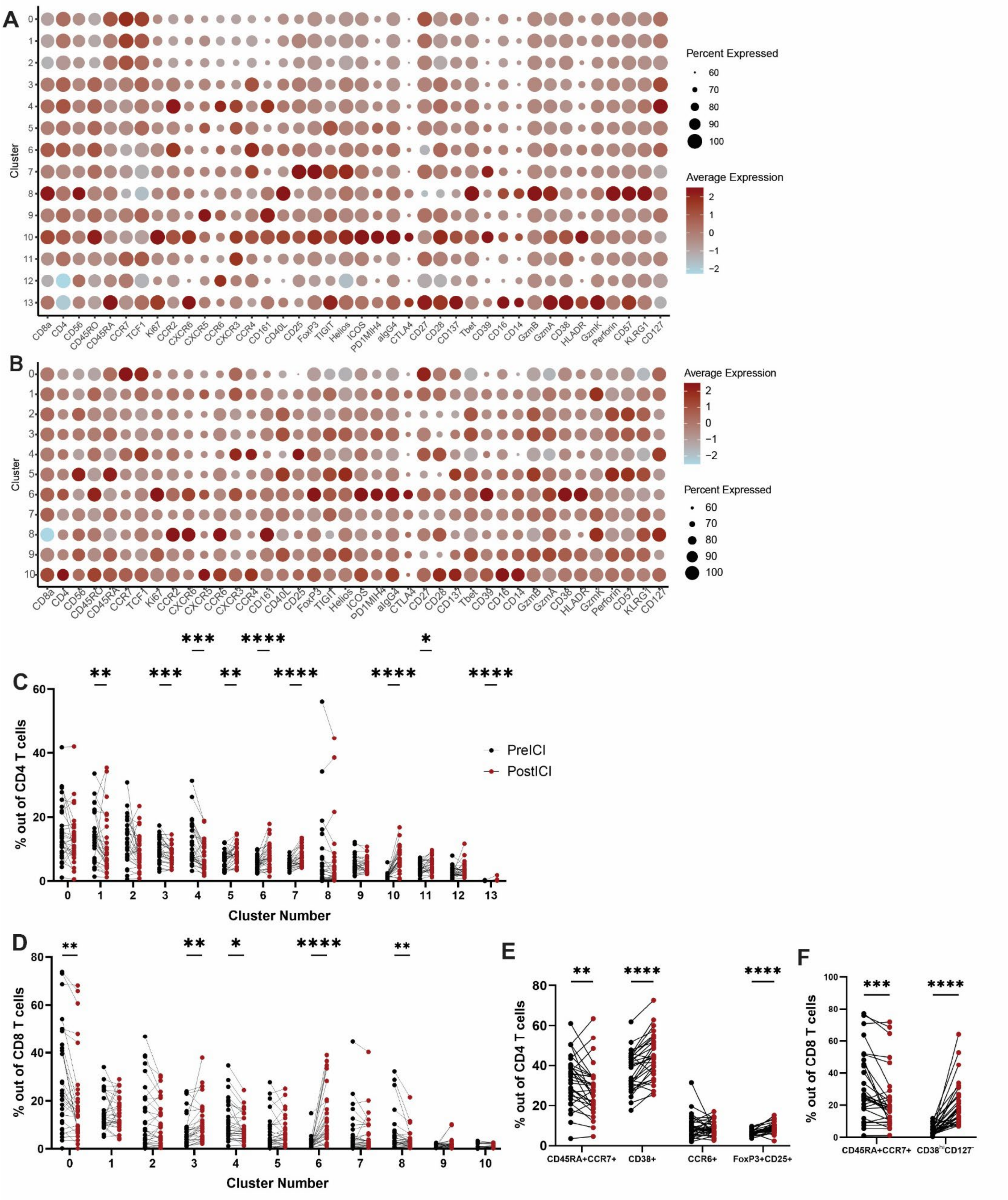
A
Cluster
0
1
2
3
4
5
6
7
8
9
10
11
12
13
CD8a
CD4
CD56
CD45RO
CD45RA
CCR7
TCF1
Ki67
CCR2
CXCR6
CXCR5
CCR6
CXCR3
CCR4
CD161
CD40L
CD25
FoxP3
TIGIT
Helios
ICOS
PD1MIH4
aIgG4
CTLA4
CD27
CD28
CD137
Tbet
CD39
CD16
CD14
GzmB
GzmA
CD38
HLADR
GzmK
Perforin
CD57
KLRG1
CD127
Percent Expressed
60
70
80
90
100
Average Expression
2
1
0
−1
−2
B
Cluster
0
1
2
3
4
5
6
7
8
9
10
CD8a
CD4
CD56
CD45RO
CD45RA
CCR7
TCF1
Ki67
CCR2
CXCR6
CXCR5
CCR6
CXCR3
CCR4
CD161
CD40L
CD25
FoxP3
TIGIT
Helios
ICOS
PD1MIH4
aIgG4
CTLA4
CD27
CD28
CD137
Tbet
CD39
CD16
CD14
GzmB
GzmA
CD38
HLADR
GzmK
Perforin
CD57
KLRG1
CD127
Average Expression
2
1
0
−1
−2
Percent Expressed
60
70
80
90
100
C
% out of CD4 T cells
0
20
40
60
PreICI
PostICI
**
***
***
**
****
****
****
*
****
0
1
2
3
4
5
6
7
8
9
10
11
12
13
Cluster Number
D
% out of CD8 T cells
0
20
40
60
80
**
**
*
****
**
0
1
2
3
4
5
6
7
8
9
10
Cluster Number
E
% out of CD4 T cells
0
20
40
60
80
**
****
****
CD45RA+CCR7+
CD38+
CCR6+
FoxP3+CD25+
F
% out of CD8 T cells
0
20
40
60
80
100
***
****
CD45RA+CCR7+
CD38hiCD127−

**Supplementary Figure 3**

A) DotPlot representation of all mass cytometry markers in CD4+ T cells subset from PBMCs (see Figure 2A-C). B) Dot Plot representation of all mass cytometry markers in CD8+ T cells subset from PBMCs (see Figure 2D-F). C) Paired analysis of frequency of all CD4 T cell and D) CD8 T cell clusters before vs after treatment per patient. test. E) Paired analysis of CD4 and F) CD8 T cells analyzed by biaxial gating. * p<0.05, **p<0.01, ***p<0.001, ****p<0.0001 by Wilcoxon paired analysis.

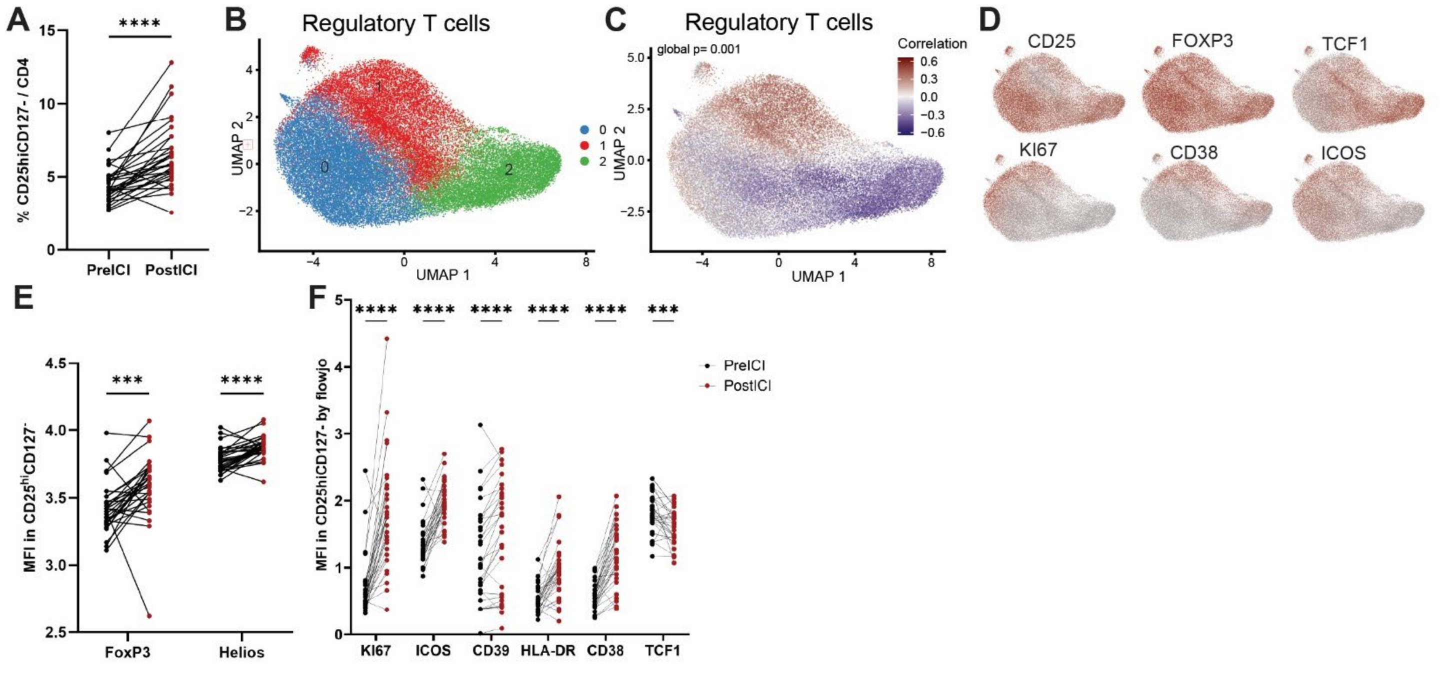


**Supplementary Figure 4** A) Paired analysis of frequency of $CD25^{hi}CD127^{-}$ CD4 T cells before and after ICI therapy by biaxial gating. B) UMAP representation following unsupervised clustering of CD4 regulatory T cells. C) CNA analysis of before treatment samples vs after treatment samples. Global p value = 0.001. D) Feature plots of the indicated markers on the Treg UMAP. E) Paired analysis of Median Fluorescent Intensity of FoxP3 and Helios or F) Ki67, ICOS, CD39, HLA-DR, and CD38 in $CD25^{hi}CD127^{-}$ CD4 T cells before and after ICI therapy. ***p<0.001, ****p<0.0001 by Wilcoxon paired analysis.

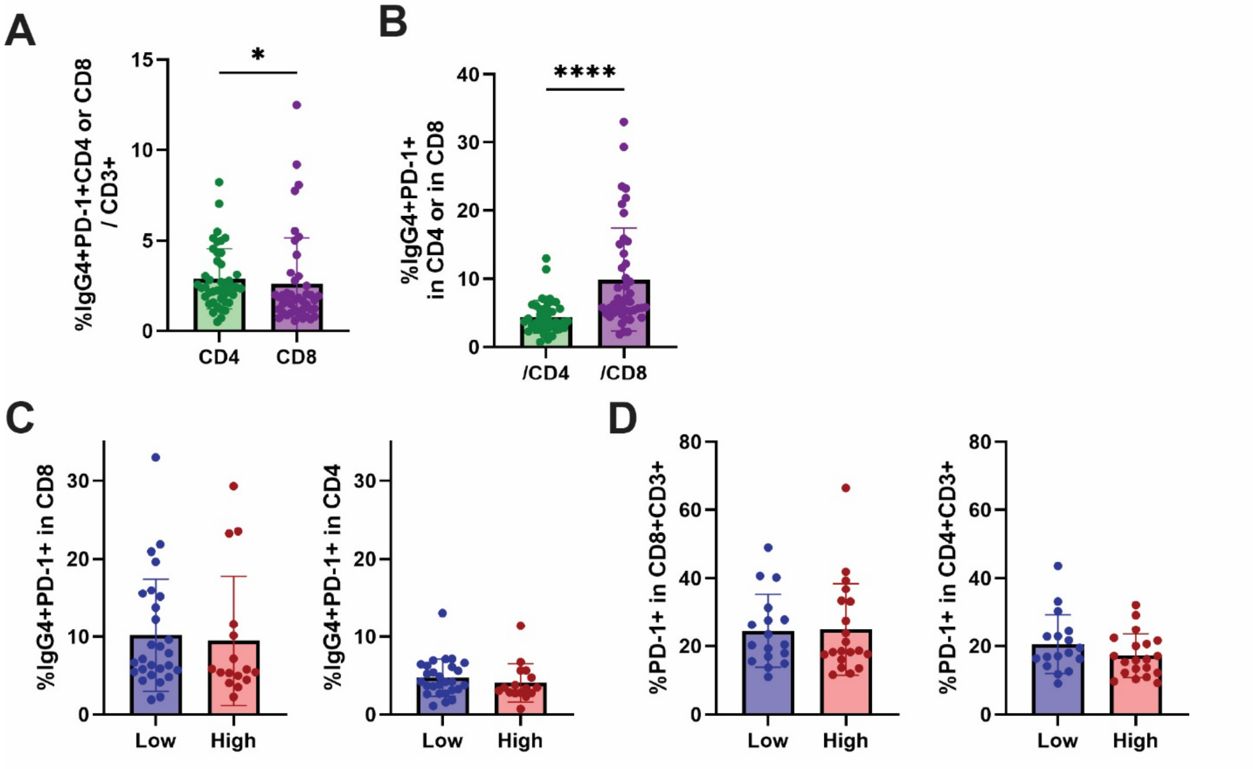


**Supplementary Figure 5**

A) Frequency of IgG4$^+$PD-1$^+$ CD4$^+$ or CD8$^+$ T cells out of CD3$^+$ T cells by gating of mass cytometry analysis in the post-treatment samples. B) Frequency of IgG4$^+$PD-1$^+$ out of CD4$^+$ or CD8$^+$ T cells by gating of mass cytometry analysis in the post-treatment samples. C) Frequency of IgG4$^+$PD-1$^+$ out of CD4$^+$ (right) or CD8$^+$ (left) T cells in patients with low or high severity of irAE by gating of mass cytometry analysis. D) Frequency of total PD-1$^+$ out of CD4$^+$ (right) or CD8$^+$ (left) T cells in pre-treatment samples from patients with low or high severity of irAE by gating of mass cytometry analysis.

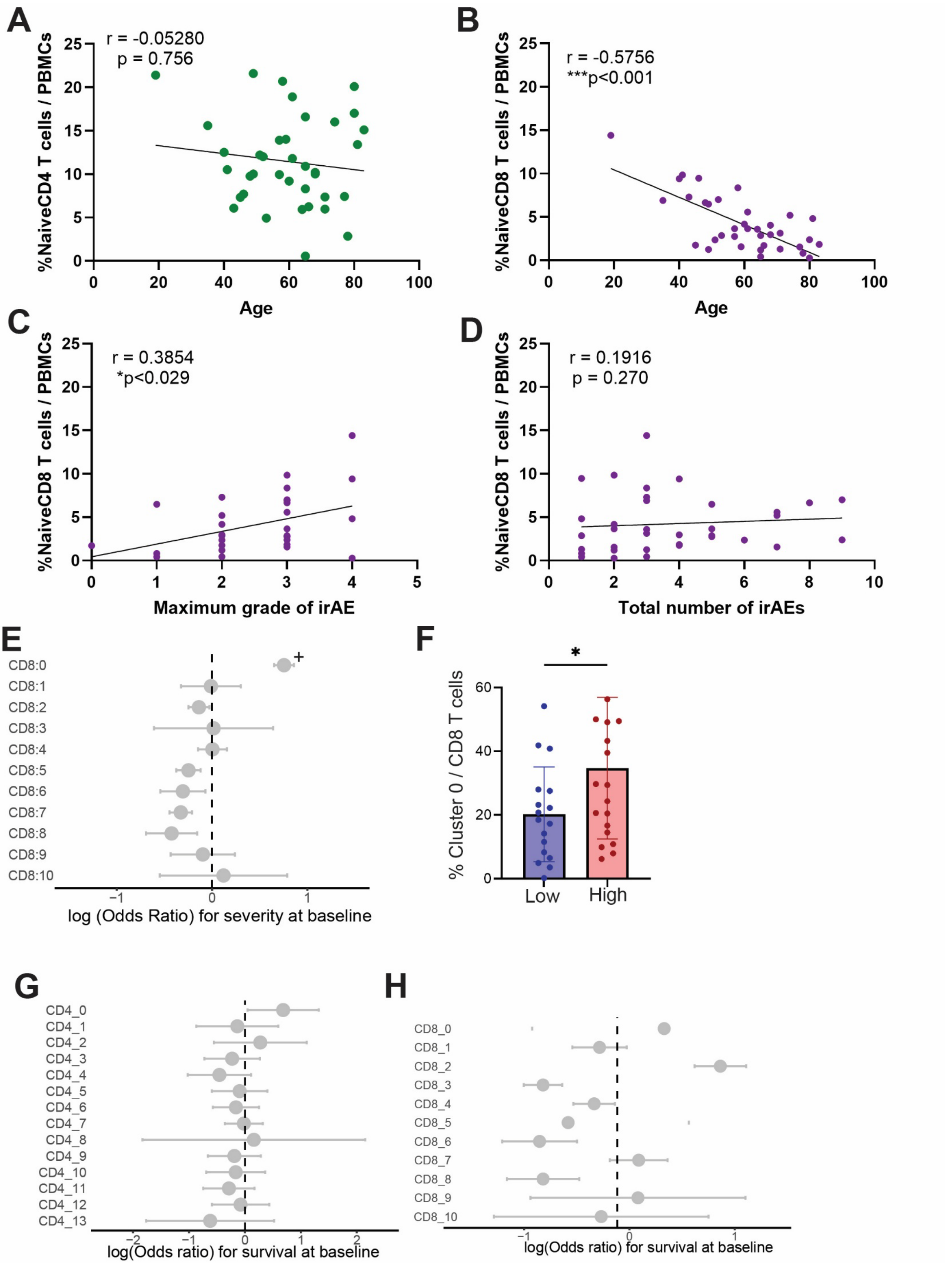

A
r = -0.05280
p = 0.756
%NaiveCD4 T cells / PBMCs
Age
B
r = -0.5756
***p<0.001
%NaiveCD8 T cells / PBMCs
Age
C
r = 0.3854
*p<0.029
%NaiveCD8 T cells / PBMCs
Maximum grade of irAE
D
r = 0.1916
p = 0.270
%NaiveCD8 T cells / PBMCs
Total number of irAEs
E
CD8:0
CD8:1
CD8:2
CD8:3
CD8:4
CD8:5
CD8:6
CD8:7
CD8:8
CD8:9
CD8:10
log (Odds Ratio) for severity at baseline
F
% Cluster 0 / CD8 T cells
Low
High
G
CD4_0
CD4_1
CD4_2
CD4_3
CD4_4
CD4_5
CD4_6
CD4_7
CD4_8
CD4_9
CD4_10
CD4_11
CD4_12
CD4_13
log(Odds ratio) for survival at baseline
H
CD8_0
CD8_1
CD8_2
CD8_3
CD8_4
CD8_5
CD8_6
CD8_7
CD8_8
CD8_9
CD8_10
log(Odds ratio) for survival at baseline

**Supplementary Figure 6**

A) Spearman correlation of proportion of gated naïve CD4+ T cells ($CD19^{-}CD14^{-}CD3^{+}CD4^{+}CD45RA^{+}CCR7^{+}$) with age. B) Spearman correlation of proportion of gated naïve CD8+ T cells ($CD19^{-}CD14^{-}CD3^{+}CD8^{+}CD45RA^{+}CCR7^{+}$) with age, C) maximum grade or irAE, and D) total number of irAEs diagnosed. E) MASC analysis of CD8 T cell clusters (see Figure 2). + indicates $p < .05$, adj $p = 0.5$ . F) Proportion of cluster 0 out of CD8 T cells. $p<0.05$ by Mann Whitney test. G) MASC analysis for overall survival in $CD4^{+}$ and H) $CD8^{+}$ T cells before treatment (survived n = 5, not survived n = 29).
